# Sterols regulate ciliary membrane dynamics and hedgehog signaling in health and disease

**DOI:** 10.1101/2025.07.15.664899

**Authors:** Antonin Lamazière, Lucie Kozlowski, Jesus Ayala-Sanmartin, Manal Farhat, Nabil Belaid, Alexandre Certain, Bertrand Isidor, Thomas Besnard, Germain Trugnan, Gregory J Pazour, Gaëtan Després, Céline Deseille, Sarah Moreno, Thibaut Eguether

## Abstract

The primary cilium is a specialized signaling hub whose function depends on a tightly regulated membrane composition. While its protein content is well-characterized, its lipid identity, particularly regarding sterols, remains poorly defined. Here, we used mass spectrometry-based lipidomics to map the sterol profile of isolated primary cilia from MDCK cells. We found that ciliary membranes are enriched in cholesterol and desmosterol while excluding precursors like 7-lathosterol and limiting others, suggesting a selective sterol barrier. Inhibiting cholesterol biosynthesis at distinct enzymatic steps led to sterol accumulation, altered ciliary membrane fluidity, and impaired Hedgehog signaling, including defective Smoothened (Smo) retention— even in the presence of a constitutively active form of Smo. These findings link sterol homeostasis to ciliary membrane properties and signaling fidelity. Our work provides a molecular framework for understanding Hedgehog-related phenotypes in disorders like Smith-Lemli-Opitz Syndrome, highlighting the importance of membrane lipid composition in developmental signaling.

## Introduction

Primary cilia are complex microtubule-based hair-shaped sensory organelles that are crucial regulators of cell signaling. The receptors and downstream effectors of many signaling proteins are enriched and sequestered in the cilium and this compartmentalization enables fine temporal and spatial regulation of pathway activation and downstream signal propagation^1^. The major cilia-linked pathway in vertebrates is Sonic Hedgehog (Shh), which plays fundamental roles during development and in adult tissue homeostasis. All the key components of this pathway are enriched in the cilium^2–5^ and their localization changes dynamically in response to its activation. When not activated, patched-1 (Ptch1), the Hedgehog ligand receptor, accumulates in the primary cilium and prevents ciliary accumulation and the activation of smoothened (Smo). Ptch1 is removed from the cilium upon ligand binding, and, consequently, Smo is derepressed and accumulates. Activated Smo subsequently activates downstream signaling, which results in the accumulation of the Gli (Glioma-associated oncogene) transcription factors at the cilium tip before their modification and translocation to the nucleus where they modulate target genes. Defects in ciliary structure and/or function lead to a set of diseases referred to as ciliopathies, a pleiotropic group of syndromes of various molecular origins (see review in^6–9^). Because Hedgehog signaling is tightly linked to the primary cilium, Hedgehog defects contribute to the phenotype of many ciliopathies causing specific symptoms encompassing craniofacial abnormalities, holoprosencephaly, syndactyly, polydactyly and heart defects. Smith-Lemli-Opitz Syndrome (SLOS, OMIM 270400), a cholesterol metabolism inborn error, share several of these Hedgehog-linked symptoms^10,11^ raising intriguing questions about the relationship between cholesterol metabolism and primary cilia in the context of Shh signaling. Cholesterol metabolism is complex, and its biosynthesis involves numerous enzymes that can give rise to various pathologies when malfunctioning^10,11^. Some of these pathologies, such as SLOS are neurodevelopmental syndromes with clear overlap with Hedgehog defects^12–14^, others, like X-linked Dominant Chondrodysplasia Punctata 2 (CDPX2 or Conradi-Hünermann-Happle syndrome, OMIM 302960), are more ambiguous and incompletely characterized in terms of mechanism with no clear links to Hedgehog signaling reported in the literature^11,15^. SLOS results from inactivating mutations in 7-dehydrocholesterol reductase (DHCR7)^16,17^, the final, rate-limiting enzyme of the cholesterol biosynthesis pathway. The enzyme blockade resulting from DHCR7 mutation leads to a decrease in the cholesterol plasma concentration and an increase in the upstream metabolites, 7-dehydrocholesterol (7-DHC) and 8-dehydrocholesterol (8-DHC)^18,19^. CDPX2 results from inactivating mutations in Δ7-Δ8 sterol isomerase (EBP), leading to lower cholesterol concentrations and higher 8-DHC and zymosterol concentrations. Since two conditions resulting in cholesterol synthesis defects present widely different outcome in terms of Shh defects, the relationship between primary cilia and cholesterol^20^ in the context of Shh signaling needs to be refined. Shh signaling has been known to be tightly linked to cholesterol for several decades as the secreted factor Shh is modified with a cholesterol moiety^21,22^. It is also well known that cholesterol and oxysterols can bind and activate Smo^23–25^. Moreover, several recent studies have presented interesting models of how ciliary cholesterol accessibility might be a major element that allow cholesterol interaction with signaling components to regulate pathway activation^26,27^. However, most studies overlook the fact that cholesterol is not the only sterol present in biological membranes and, consequently, the impact of cholesterol metabolites other than oxysterols (e.g., lanosterol, 7DHC, 8DHC, and desmosterol) on the membrane biophysical properties and subsequent signaling has not been studied to date. Hence, we designed a study to describe the sterolome (i.e. quantitative sterol composition) of the primary ciliary membrane in mammalian cells as well as in situations mimicking the defects in SLOS and CDPX2 patients. To do so, we used a multidisciplinary approach comprising mass spectrometry-based metabolomics of isolated mammalian primary cilia, biophysics of model membranes and functional experiment of Hedgehog signaling in patients and CRISPR-generated cells.

Our findings demonstrate that the primary ciliary membrane contains a subset of sterols in specific proportions whose homeostasis is tightly regulated by the cell. Modulation of the cholesterol synthesis pathway led to drastic changes in the sterol composition of cell membranes, including the ciliary membrane, although primary cilia exhibited some resistance to these changes. Indeed, under pharmacological treatment mimicking the SLOS or CDPX2 enzyme blockades, the gross variations in the primary ciliary membrane were comparable to what happens in other cell membranes, but with differing intensity, further highlighting the tight regulation of primary ciliary membrane homeostasis. Ciliary membranes were enriched in cholesterol and desmosterol; however precursors such as 7-lathosterol were lacking and others were limited, suggesting a selective sterol barrier. Disturbing that specific homeostasis led to the accumulation of sterol metabolites with membrane stiffening properties, which could be the origin of the Hedgehog defects exhibited by pharmacologically treated cells, human SLOS patients and CDPX2 CRISPR-generated fibroblasts. Indeed, cells mimicking both pathologies exhibited strong defects in of Smo behavior when chemically activated by SAG (Smoothened Agonist) and defective activation when the constitutively active protein SmoM2 was overexpressed. This finding was unexpected for CDPX2, as this pathology has not been previously associated with Hedgehog defects. Thus our study sheds new light on the relationship between primary cilia cholesterol and cholesterol metabolism defects.

## Materials and Methods

### Primary cilia isolation

Full procedure will be described in detail elsewhere. Briefly, MDCK cells were grown to confluency and serum starved during 6 days prior cilia isolation. Cells from 5 x 150 mm^2^ dishes were then washed with PBS-EDTA 0,04% for 10 min and scraped in PBS. After pelleting the cells by centrifugation, the pellet was resuspended in 0,75 mM dibucaine (Sigma Aldrich) in PHEM (Pipes 45 mM pH 6.9, Hepes 45 mM pH 6.9, EGTA 10 mM pH6.9, MgCl2 5 mM) and rotated for 15 minutes at room temperature. Taxol was then added to obtain a 2 µM final concentration and then rotated 10 minutes à room temperature. Thereafter, cilia were found in the supernatant. Supernatant was then deposited on top of a sucrose gradient (70%, 50%, 25%) and centrifuged for 1h15 at 100 000g. Cilia were collected at the interface between the 25% and 50% concentrations. Quality of the preparation was assessed by immunofluorescence with ciliary markers (Arl13b for the membrane and acetylated tubulin for the axoneme). Cilia were kept at −80°C until they were used.

### Sterol quantification by GC-MS

Sterols in cell homogenates or isolated cilia fraction were extracted with a solvent mixture containing chloroform/methanol 2/1 (v/v) spiked with internal standards [epicoprostanol, 2H7-7-lathosterol, 2H6-desmosterol, 2H6-lanosterol] (Avanti Polar Lipids). Lipids were partitioned in chloroform after the addition of saline and saponified by methanolic potassium hydroxide (0.5 N, 60 °C, 15 min). The released fatty acids were methylated with BF3-methanol (12%, 60 °C, 15 min) in order to avoid their interference with the chromatography of sterols. The sterols were re-extracted in hexane and silylated, as described previously61. The trimethylsilylether derivatives of the sterols were separated by gas chromatography (GC) (Hewlett–Packard 6890 series) in a medium polarity capillary column RTX-65, (65% diphenyl 35% dimethyl polysiloxane, length 30 m, diameter 0.32 mm, film thickness 0.25 μm (Restesk, Evry, France)). The mass spectrometer (Agilent 5973 inert XL) in series with the gas chromatography was set up for detection of positive ions. Ions were produced in the electron impact mode at 70 eV. They were identified by the fragmentogram in the scanning mode and quantified by selective monitoring of the specific ions after normalization and calibration with the appropriate internal and external standards [epicoprostanol m/z 370, 2H7-7-lathosterol m/z 465, 2H6-desmosterol m/z 358, 2H6-lanosterol m/z 504, cholesterol m/z 329, zymosterol m/z, 7-dehydrocholesterol m/z 325, 8-dehydrocholesterol m/z 325, desmosterol m/z 343, lanosterol m/z 393]

### Proteomics analysis

#### Purification

6 fractions of 3 different preparations of primary cilia isolated from MDCK cells were used. The high percentage of sucrose from the fractions was diluted 10 times with PBS and cilia were subsequently concentrated using Amicon ultra filters 10000 HMWCO (Merck Millipore, Burlington, MA, USA). The remaining 1 mL was then centrifuged at 12500g to pellet the cilia and remove the supernatant. The pellet was resuspended in 20mL denaturing gel loading buffer (Tris-HCl 125 mM pH6.8, Glycerol 20% v/v, SDS 4% v/v, β-Mercaptoethanol 10% v/v, bromophenol blue) buffer and subjected to SDS-PAGE migration in a 6% gel. The gel was sent to Umass Chan Medical school proteomics facility for shot-gun proteomics analysis on an Orbitrap Fusion Lumos Tribrid.

#### Database searching

Charge state deconvolution and deisotoping were not performed. All MS/MS samples were analyzed using Mascot (Matrix Science, London, UK; version Mascot in Proteome Discoverer 2.1.1.21). Mascot was set up to search CanisFamiliarisNCBI assuming the digestion enzyme strict trypsin. Mascot was searched with a fragment ion mass tolerance of 0,050 Da and a parent ion tolerance of 10,0 PPM. Carbamidomethyl of cysteine was specified in Mascot as a fixed modification. Gln->pyro-Glu of the n-terminus, oxidation of methionine and acetyl of the n-terminus were specified in Mascot as variable modifications.

#### Criteria for protein identification

Scaffold (version Scaffold_4.11.0, Proteome Software Inc., Portland, OR) was used to validate MS/MS based peptide and protein identifications. Peptide identifications were accepted if they could be established at greater than 90,0 % probability by the Peptide Prophet algorithm^28^ with Scaffold delta-mass correction. Protein identifications were accepted if they could be established at greater than 99,0 % probability and contained at least 2 identified peptides. Protein probabilities were assigned by the Protein Prophet algorithm^29^. Proteins that contained similar peptides and could not be differentiated based on MS/MS analysis alone were grouped to satisfy the principles of parsimony. Proteins sharing significant peptide evidence were grouped into clusters.

#### PANTHER analysis

PANTHER overrepresentation Test^38^ (Released 2024 08 07) was launched using GO Ontology database (DOI 10.5281/zenodo.15066566) Released 2025 03 16. A subset of the 3000 proteins most represented in 3 independent proteomics experiment was submitted and compared to the Canis lupus familiaris database using Fisher’s exact test, with the Benjamini–Hochberg false discovery rate (FDR) correction.

##### Large unilamellar vesicles (LUV)

LUVs of 100 nm diameter of varying lipid compositions were prepared by extrusion in PBS, as described in ^30^. Briefly, the lipids in a chloroform/methanol solution (1/1) were mixed in the desired proportions and dried. The lipids were re-suspended in PBS by strong vortexing. The resulting MLVs (multilamellar vesicles) at 0.5 mg/ml were extruded by passing them through 100 nm pore polycarbonate filters. LUVs containing the most important lipids of the outer leaflet of the plasma membrane: phosphatidylcholine (PC), sphingomyelin (SM) and cholesterol (chol) or different sterols were prepared. Egg yolk L-α-phosphatidylcholine (PC), egg yolk sphingomyelin (SM) and cholesterol (chol) were purchased from Sigma-Aldrich. Desmosterol (Desmo), 7-dehydrocholesterol (7dhc) and 8-dehydrocholesterol (8dhc) were purchased from Avanti polar lipids. Di-4-ANEPPDHQ (ANE) was obtained from Invitrogen

##### Fluorescence recording

Fluorescence measurements were performed with a Tecan infinite M200 fluorimeter. 20 µg of LUVs were suspended in 200 µl of buffer in wells of a 96 wells cell culture plate (Falcon). The probe di-4-ANNEPDHQ (ANE) was used at 1 μM. This environmental probe, like Laurdan, is able to monitor membrane fluidity. The emission spectra were recorded from 500 to 750 nm using 485 nm excitation. Spectra were recorded at temperatures of 25°C and 37°C. The excitation generalized polarization (GP) for ANE was calculated as GP = (I_570_-I_630_)/(I_570_+I_630_), where I_570_ and I_630_ are the fluorescence intensities at the maximum emission wavelength in the ordered (570 nm) and disordered (630 nm) phases respectively. The statistical significance of the GP differences was assessed by student t test with GraphPad Prism. ** P<0.01, *** P<0.001.

### Cell culture

*Murine Embryonic Fibroblasts (MEF)* and their derivatives were grown at 37°C in 5% CO_2_ in Dulbecco’s modified Eagle’s medium (DMEM; Gibco) with 10% FBS and 1% Penicillin-Streptomycin (Gibco). For SAG (Smoothened AGonist) experiments, cells were plated at near confluent densities and serum starved (same culture medium described above but with 0.25% FBS) for 48 hours prior to treatment to allow ciliation. SAG (Calbiochem) was used at 400 nM. *Madin Darby Canine Kidney cells (MDCK)* were grown at 37°C in 5% CO_2_ in Dulbecco’s modified Eagle’s medium - F12 (DMEM-F12; Gibco) with 10% FBS and 1% Penicillin-Streptomycin (Gibco). Before cilia isolation, cells were plated at near confluent densities and serum starved (same culture medium described above but with 0.25% FBS) for 6 days prior to treatment to allow ciliation.

*Human Embryonic Kidney cells (HEK293-T)* were grown at 37°C in 5% CO_2_ in Dulbecco’s modified Eagle’s medium (DMEM; Gibco) with 10% FBS and 1% Penicillin-Streptomycin (Gibco).

Presence of mycoplasma was monitored by PCR with the Venor Gem Onestep (Minerva).

#### Human patient fibroblasts

Human fibroblasts were collected from one SLOS patient and a healthy donor harboring two mutations - c.725G>A (Arg242His) and c.906C>G (Phe302Leu) - in their *DHCR7* gene (11q13.4). Patients gave full consent for the use of this biological material in research procedures.

#### Plasmids

The following plasmid was transfected into cells via lentiviral infection:

**SmoM2-mCherry** in pHAGE_DN_CMV vector (originally from D. Nedelcu and A. Salic). All MEF cell lines following infection were selected with appropriate antibiotic and dilution cloned with flow cytometry using morphological characteristics to add one cell per well in a 96-well plate.

**TE139** in LentiCRISPRv2 vector (Addgene #98290) is a plasmid designed to Knock-out the protein EBP (Emopamil Binding Protein or cholestenol delta-isomerase or Δ7-Δ8 isomerase) through lentiviral infection of Guide RNA and Cas9 protein. Guide RNA sequence (**CACCGCGACGGATTCCAGCA**) was designed using Chop-Chop (https://chopchop.cbu.uib.no/).

**TE141** in LentiCRISPRv2 vector (Addgene #98290) is a plasmid designed to Knock-out the protein EBP (Emopamil Binding Protein or cholestenol delta-isomerase or Δ7-Δ8 isomerase) through lentiviral infection of Guide RNA and Cas9 protein. Guide RNA sequence (**CTCCGTGCTGGAATCCGTCG**) was designed using Chop-Chop.

#### Pharmacological treatments

MDCK cells were plated on Day 0 in DMEM-F12 supplemented with 10% serum, and then serum starved on day 1 with DMEM-F12 supplemented with 0,25% serum to allow them to form cilia. At day 6, cells were treated for 48h with either 20 µM AY9944 (Sigma-Aldrich), 4 µM Tamoxifen (Fluka) or 5 µM Simvastatin (Sigma-Aldrich). At day 7, cilia were harvested to perform mass spectrometry experiments.

MEF cells were plated on Day 0 in DMEM supplemented with 10% serum, and then serum starved on Day 1 with DMEM supplemented with 0,25% serum to allow them to form cilia. At Day 3, cells were treated for 48h with either 5 µM AY9944 (Sigma-Aldrich), 2,5 µM Tamoxifen (Fluka) or 2,5 µM Simvastatin (Sigma-Aldrich). At day 4, cilia were harvested to perform qPCR or IF experiments.

### Lentivirus production

The pHAGE system is a third generation lentiviral system comprising four distinct packaging vectors (Tat, Rev, Gag/Pol, VSV-g). Packaging of the virus was performed using HEK 293T cells as a host for co-transfection of the backbone vector and the four packaging vectors in the following proportion [Backbone: 5; Tat: 0.5; Rev: 0.5; Gag/Pol: 0.5; VSV-g: 1]. Effectene (Qiagen) or Calcium Chloride were used as transfection reagent used was. After 48 hours of virus formation, supernatant was harvested and filtered through a 0.45 µm filter. Lenti-x concentrator (Clontech) was added to the filtrate following manufacturer’s protocol. After centrifugation, pellet was resuspended in DMEM and polybrene was added at a final concentration of 5 g/mL. Viruses were finally added overnight to the plate containing cells to infect around 10-20% confluence. Cells were selected with the appropriate antibiotic depending on the plasmid used.

### mRNA Analysis

Cells were kept frozen at −80°C in RNA Lysis Buffer (RLT) supplemented with β-mercaptoethanol as per manufacturer protocol until extraction (RNeasy, Qiagen). RNA was extracted using QiaShredder columns followed by RNeasy kits with on-column DNA digestion (Qiagen). RNA concentration was determined spectrophotometrically. cDNA was synthesized using LunaScript (New England Biolabs). Thermal cycling was performed in either a StepOne Plus system (Applied Biosystems) or a CFX Opus96 (BioRad). Both apparatuses showed comparable performances. Specific primers sequences are described in Table 1. Primers were designed with the Primer3 web-based software (https://bioinfo.ut.ee/primer3/). Fold changes in gene expression were calculated after normalizing data to GAPDH expression and then normalized to untreated controls (ΔΔCt). PCR was performed using 15 μl reactions containing 2.5 ng of first strand cDNA, 0.5 μM forward and reverse primers, and PowerSYBR green PCR master mix (Applied Biosystems). qPCR reactions were performed in triplicate using the following protocol: 50 °C for 2 min, 95 °C for 10 min, 40 cycles at 95 °C for 15 s, and 60 °C for 60 s, then dissociated to verify a single amplicon. Quantitative RT-PCR primers are described in table 3.

**Table 1.**
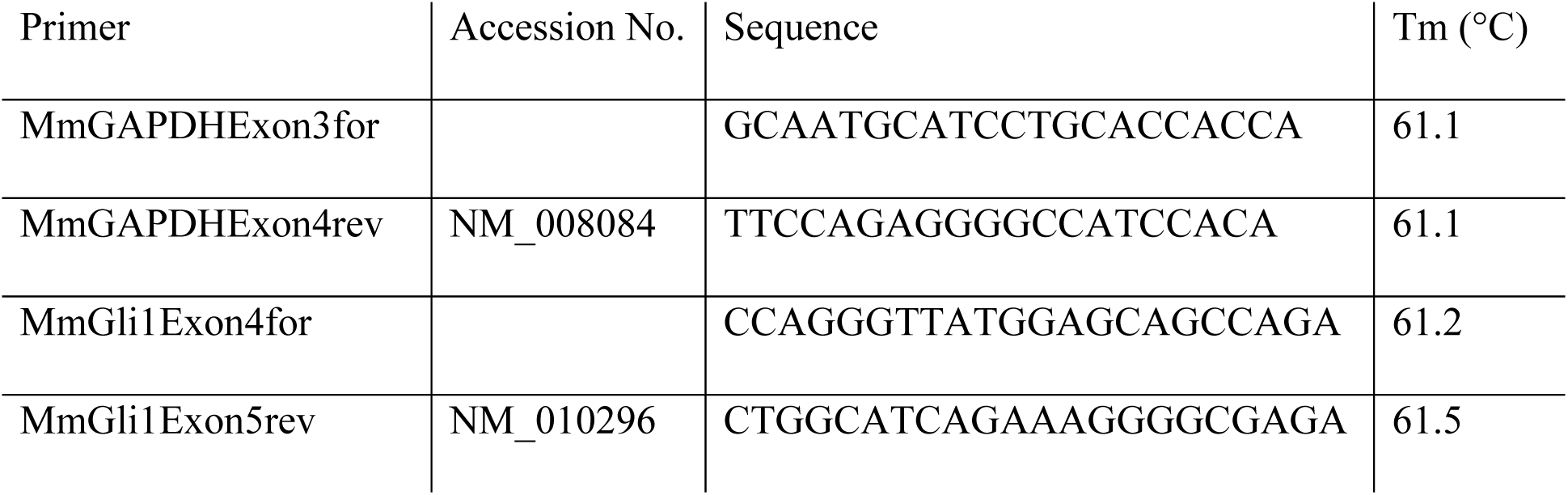
Quantitative RT-PCR primers.

### Immunofluorescence

Cells were fixed with 2% paraformaldehyde for 15 min, permeabilized with 0.1% Triton-X-100 for 5 min and treated with 0.05% SDS for 5 min to retrieve antigens. The primary antibodies are described in table 2.

**Table 2.**
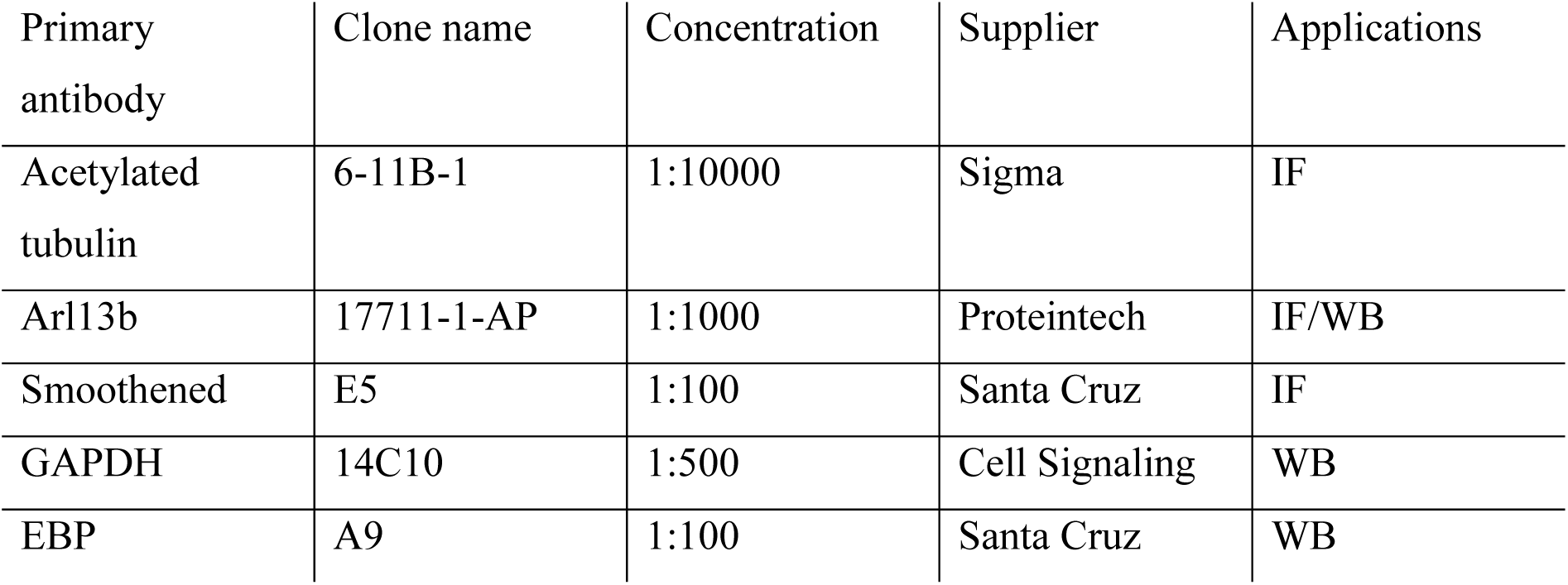
Antibodies.

### Quantification of fluorescence in cilia

Pictures of cells harboring primary cilia were taken randomly with an Apotome Microscope (Zeiss) with Z-stacks. After Z-projection of the stack in a single image, cilia were counted manually and analyzed with the tools integrated to ImageJ (Rasband, W.S., ImageJ, U. S. National Institutes of Health, Bethesda, Maryland, USA, https://imagej.nih.gov/ij/, 1997-2016.) to extract the percentage of ciliated cells in a field, length, and fluorescence intensity. For quantification of Smo, the average Smo fluorescence intensity was measured within a variable area comprising total detectable Smo staining in the primary cilium. For each measurement, background was also quantified in the cytoplasm and subtracted from mean Smo intensity.

### Protein Analysis

For western blots, MEFs were lysed directly into denaturing gel loading buffer (Tris-HCl 125 mM pH6.8, Glycerol 20% v/v, SDS 4% v/v, β-Mercaptoethanol 10% v/v, bromophenol blue) and sonicated (20% energy). Western blots were developed by chemiluminescence (Super Signal West Pico plus, Pierce Thermo) and imaged using a iBright 1500 (Invitrogen, ThermoFisher).

### Statistics

Data groups were compared using non-parametric Mann-Whitney or Kolmogorov-Smirnov tests (two groups) or one-way Kruskall-Wallis test (more than two groups) followed by Turkey’s or Dunett’s multiple comparison tests. Parametric one-way Anova followed by Holms-Sidak’s multiple comparison test were also used after verifying the normality of the distribution by the Shapiro-Wilk test. All analyses were made using the GraphPad Prism 10 software. Differences between groups were considered statistically significant if p < 0.05. Statistical significance is denoted with asterisks (*p=0.01 – 0.05; **p=0.001-0.01; ***p=0.0001-0,001: ****p<0,0001). Error bars are all S.D. Center values are all averages.

## Results

### Mammalian primary cilia have a unique sterol composition

Numerous studies have indicated that the protein and lipid composition of primary cilia is unique^31–33^. The evidence for a specific protein composition in mammalian cells is overwhelming but, however, studies on ciliary lipid composition, regulation, and function have lagged behind, partly due to the lack of efficient primary cilia isolation methods. To gain deeper insight into this crucial question, we established a method to purify Madin-Darby canine kidney (MDCK) cells primary cilia via the use of dibucaine, a local anesthetic that acts as a Na-channel blocker, and can deciliate MDCK cells in tissue culture (Fig.1a). Immunofluorescence with the widely used ciliary markers Arl13b (membrane) and acetylated tubulin (axoneme) (Fig.1b) revealed that this method allows efficient recovery of cilia of various lengths confirmed by Arl13B enrichment as shown by western blotting (Fig.1c) of a purified ciliary fraction compared with that of whole cell extract prepared with deciliated cells. To evaluate the degree of enrichment of primary cilia, we performed mass spectrometry-based proteomics on the ciliary fraction and recovered 210 of the ciliary-specific proteins from the Cilia Carta database^34^ with at least 3 peptides in 3 independent experiments. The dataset included 18 IFT proteins, 4 BBS proteins, several components of the Hedgehog pathway (e.g., SuFu and Arl13b), axonemal proteins (Tubulin, Kinesin and Dynein chains) and centrosomal proteins (Centrin, CEPs, Gamma Tub and associated proteins) indicating that the basal body might be partially recovered. The dataset demonstrates a 70% overlap with the ciliary proteome published by Ishikawa and collegues^35^, and a 41% overlap with the CysCilia consortium potential ciliary proteins list^36^. A gene ontology network created with these 210 ciliary genes is presented in Figure 1d, further illustrating the diversity of the ciliary compartment from which the proteins were retrieved. To better describe the dataset with regard to enrichment in cilia versus other contaminant organelles, we performed a Gene Ontology Enrichment Analysis using the PANTHER database^38^. This tool analyses the overrepresentation of the proteins by comparing the background frequency, i.e., the number of genes annotated to a GO term in the considered genome (here Canis lupus familiaris) to the sample frequency, the number of genes/proteins annotated to that GO term in the input list. Following this analysis, several key ciliary components were present in the overrepresented list with good calculated p values (FDR : false discovery rate) (see Table 3): ciliary inversin compartment, intraciliary transport particle B, BBSome, intraciliary transport particle, ciliary base, ciliary basal body, centriolar satellite, microtubule associated complex, microtubule, centrosome, microtubule organizing center and cilium. Some GO terms associated with the overrepresented proteins were not exclusively related to primary cilia, but have been linked to ciliary function in recent studies, for example, nuclear pore, p-body, translation initiation factors^40^, exocyst complex^42^ and AP-1 adaptor complexe^44^. Others demonstrated no clear link to cilia biology, thus these components were probable contaminants of the cilia preparation, such as mitochondria, actin cytoskeleton or ribosomes. Finally, some GO terms such as axon-related component were difficult to interpret since the isolate originated from kidney cells. These findings illustrate the difficulty of this type of approach, which is highly dependent on the quality of the proteins and genome annotation, which is probably suboptimal for *Canis lupus familiaris*. Taken together, the results show that cilia isolation allows efficient retrieval of mammalian primary cilia with good enrichment of the ciliary fraction, but, as expected, leads to contaminants from other cell compartments.

**Figure 1.**
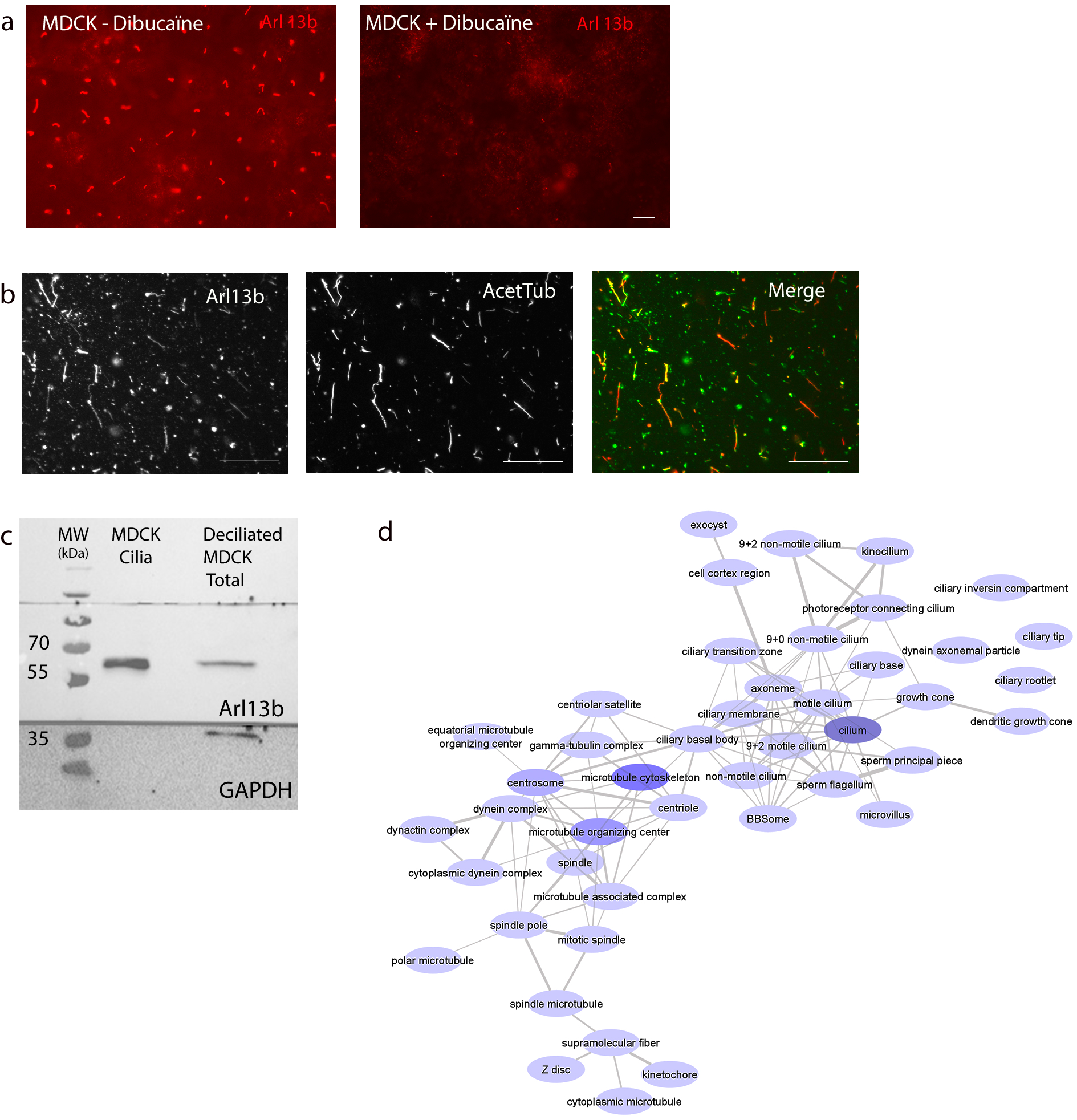
Primary cilia isolation with dibucaine in MDCK cells. (a) Effect of dibucaine on primary cilia of MDCK cells. Immunofluorescence with Arl13b antibody on MDCK cells serum starved for 6 days with or without dibucaine addition at day 6. Scale bar : 10 µm (b) Immunofluorescence with Arl13b (ciliary membrane) and acetylated tubulin (AcetTub, axoneme) antibodies on purified cilia after sucrose gradient. Merge shows Arl13b in Green and Acetylated Tubulin in red. Scale bar : 10 µm (c) Western Blotting with Arl13b and GAPDH antibodies showing enrichment of Arl13b in the ciliary fraction (MDCK Cilia) compared to the rest of the cell after deciliation (Deciliated MDCK total). (d) GO term analysis of the proteomic data showing that retrieved peptides span most compartments of the cilium.

**Table 3.** Overrepresented ciliary GO terms after PANTHER analysis.

| GO cellular component complete | Canis lupus familiaris (20969) | Protein Set (2376) | Fold enrichment | Raw P-value | FDR |
| --- | --- | --- | --- | --- | --- |
| ciliary inversin compartment | 3 | 3 | 8,83 | 1,45E-03 | 1,04E-02 |
| intraciliary transport particle B | 17 | 13 | 6,75 | 7,54E-10 | 1,60E-08 |
| BBSome | 8 | 6 | 6,62 | 4,81E-05 | 4,52E-04 |
| intraciliary transport particle | 26 | 16 | 5,43 | 1,23E-09 | 2,55E-08 |
| ciliary base | 32 | 12 | 3,31 | 1,11E-04 | 1,01E-03 |
| ciliary basal body | 246 | 58 | 2,08 | 4,40E-08 | 6,47E-07 |
| centriolar satellite | 100 | 23 | 2,03 | 7,54E-04 | 5,76E-03 |
| microtubule associated complex | 129 | 27 | 1,85 | 1,83E-03 | 1,28E-02 |
| microtubule | 364 | 74 | 1,79 | 6,14E-07 | 8,28E-06 |
| centrosome | 519 | 103 | 1,75 | 1,38E-08 | 2,15E-07 |
| microtubule organizing center | 696 | 132 | 1,67 | 1,86E-09 | 3,58E-08 |
| cilium | 714 | 109 | 1,35 | 1,15E-03 | 8,43E-03 |

Purified ciliary fractions were then subjected to mass spectrometry-based lipidomics to quantify the various ciliary membrane sterols species. This series of experiments revealed that the sterol composition of the ciliary membrane differs from that of the other cell membranes (Fig.2). Three sterols are present at relative higher concentrations in the ciliary membrane than in the cell membrane: cholesterol, with an average 2.5-fold increase; desmosterol, with a 2-fold increase and lanosterol, with an increasing trend but lack of statistical significance due to its low levels and greater variability. Conversely, two metabolites were present at a lower relative concentration in the ciliary membrane: 7-lathosterol, which, as we demonstrated throughout this manuscript, is always excluded from the ciliary membrane, and 8-dehydrocholesterol (8-DHC) whose concentration is 5-fold lower than that of the cell membranes. 7-DHC appears to be present in the same proportion in both compartments. Taken together, these results show that the sterol content of the mammalian primary cilia membrane is closely regulated.

**Figure 2.**
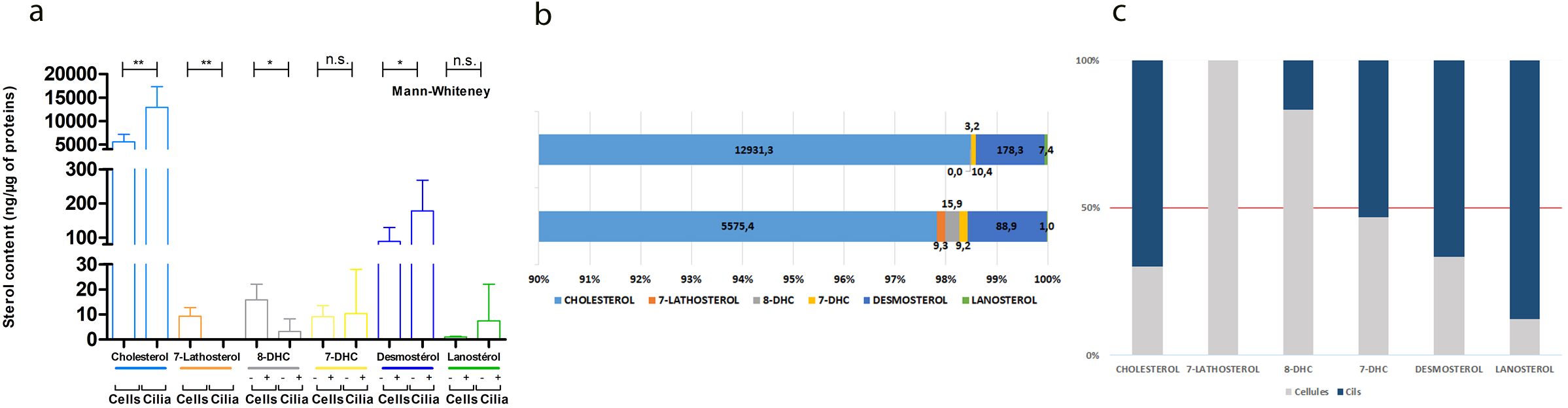
Sterolome of a mammalian primary cilium. (a) Comparison of the sterol mass spectrometry results from 6 independent primary cilia isolates (Cilia) and the rest of the cell membranes (Cells). (b) Proportion of the different sterols in isolated cilia and the other cell membranes. (c) Combined proportion of the different sterols in isolated cilia (Cilia, in blue) and the other cell membranes (Cells, in grey). The 50% limit is when, in proportion, there is the same amount of a specific sterol species in the cilium and in the other cell membranes. Above, the considered species is enriched in the cilium, below, it is enriched in the cell membranes. Sterols quantified in all panels : Cholesterol, 7-Lathosterol, 8-Dehydrocholesterol (8-DHC), 7-Dehydrocholesterol (7-DHC), Desmosterol and Lanosterol. n.s : not significant * P<0.05 ** P<0.01, Mann-Whiteney Test.

### Sterol composition modulation has different effects on the ciliary membrane than on the other cell membranes

To better understand whether a change in sterol composition of the ciliary membrane could explain the Hedgehog defects in SLOS patients, we modulated the cholesterol synthesis pathway using pharmacological agents. We designed three assays to mimic different pathophysiological situations. First, we used AY9944, a well-known DHCR7 blocker that recapitulates the enzyme defect found in SLOS patients^37^. Then, using tamoxifen, we targeted the Δ7-Δ8 isomerase also known as emopamil binding protein (EBP)^39,41^, which blockade is responsible for the X-linked chondrodysplasia punctata type 2 disease in human (CDPX2), a disease in which patients do not usually exhibit Hedgehog-linked symptoms. Finally, as DHCR7 and Δ7-Δ8 isomerase control downstream steps of cholesterol synthesis, we included a treatment with simvastatin, a member of the widely prescribed family of cholesterol-lowering drugs, known as statins, which targets the enzyme HMGCoA reductase, one of the upstream key enzyme of the cholesterol synthesis pathway.

After the treatments (see, Fig.3a and methods section), we examined the composition of the ciliary membrane by mass spectrometry and compared it to that of the fraction containing the other membranes. All treatments led to expected variations in the global cell membrane fraction considering the enzyme targeted by the different drugs.

**Figure 3.**
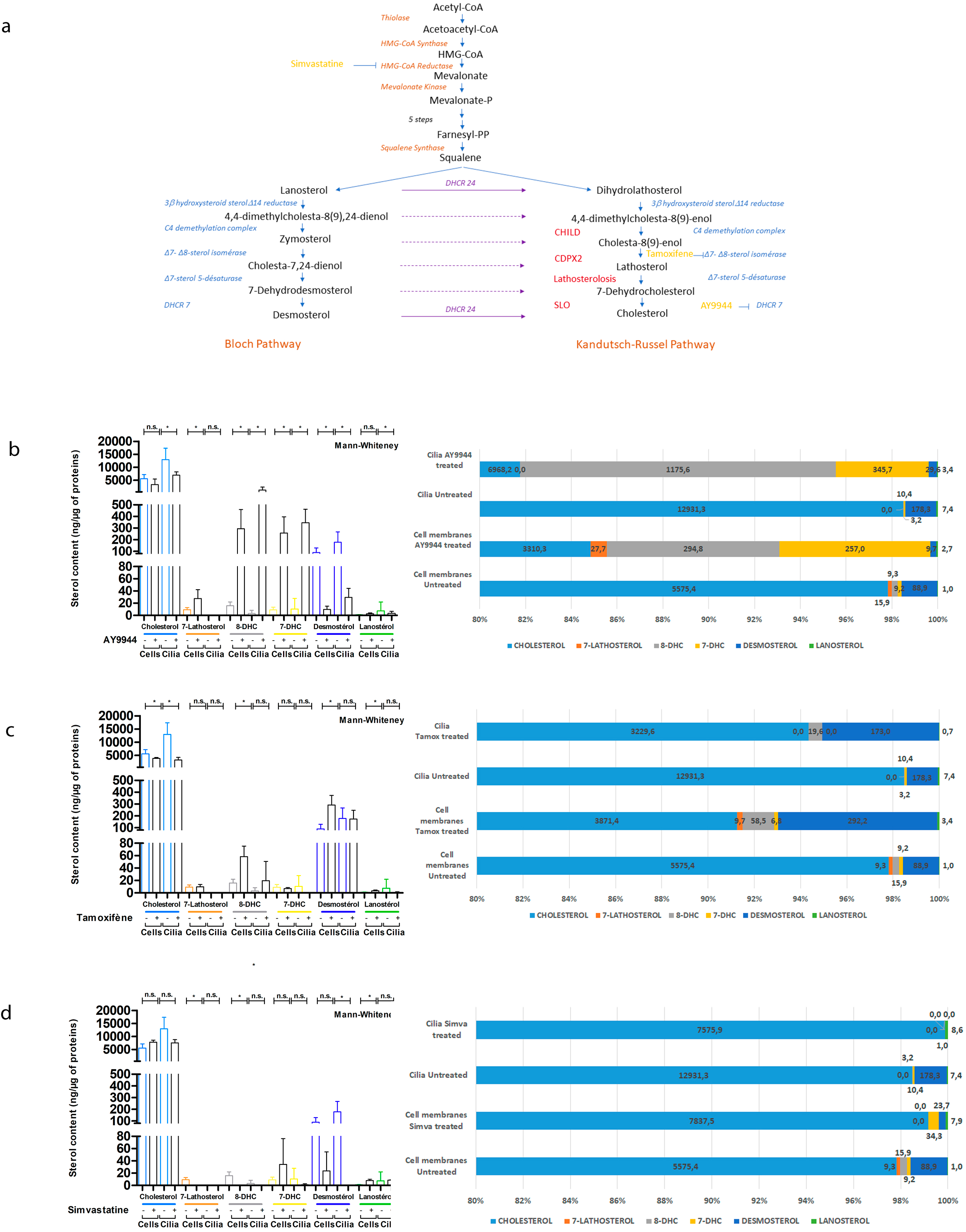
Sterolome of membranes after treatments mimicking the SLOS and CDPX2 defects. (a) Cholesterol synthesis pathways. Orange: pre-squalene enzymes, Blue: post-squalene enzymes, Red: human pathologies associated with mutations in the corresponding enzyme, Yellow: enzyme inhibitors. Purple: enzymes converting sterol species from the Bloch pathway to the Kandutsch-Russel pathway. Comparisons of absolute quantification (Left panels) and percentage (Right panels) of the sterol mass spectrometry results from primary cilia isolates (Cilia) and the rest of the cell membranes (Cells) with or without (+, -) AY9944 treatment (b), Tamoxifen treatment (c) and Simvastatin treatment (d). Sterols quantified : Cholesterol, 7-Lathosterol, 8-Dehydrocholesterol (8-DHC), 7-Dehydrocholesterol (7-DHC), Desmosterol and Lanosterol. Results from three independent experiments. n.s : not significant, * P<0.05, Mann-Whiteney Test. Errors bars sow Means ± SD.

After AY9944 treatment (Fig.3b) we observed an accumulation of sterol metabolites produced upstream of the DHCR7 blockade, namely 7-Lathosterol, 7-DHC, and 8-DHC, as well as a decrease in metabolites produced downstream of the blockade, such as desmosterol and cholesterol. Notably, AY9944 treatment led to a greater reduction in cholesterol content as well as a greater increase in 7-DHC and 8-DHC in cilia than in the other cell membranes. Moreover, it is striking that although DHCR7 blockade results in increased 7-Lathosterol concentration in cell membranes, this metabolite remained excluded from the ciliary membrane.

In cell membranes, tamoxifen treatment (Fig.3c) resulted in the expected decrease in cholesterol concentration as well as an increase in 8-DHC and desmosterol concentrations. In contrast, within the ciliary membrane, the decrease in cholesterol was more pronounced, whereas the increase in 8-DHC was smaller and didn’t reach statistical significance due to experimental variability. Finally, no increase in desmosterol was observed in the ciliary membrane.

Simvastatin treatment (Fig.3d) resulted in only subtle changes, with a moderate decrease in cholesterol, 7-lathosterol, 8-DHC, and desmosterol as well as a modest increase in lanosterol. The most marked change was observed in the ciliary compartment, where desmosterol was excluded from the ciliary membrane after treatment.

Taken together, these results show that the homeostasis of the primary ciliary membrane is tightly regulated, and selectively restricts membrane access to certain metabolites, somewhat compensating for or exacerbating the drug’s action depending on the considered metabolite.

### Membrane-stiffening sterols accumulate after DHCR7 blockade in model membranes

To gain more insight into the effects of sterol composition on membrane lipid compaction (fluidity) we performed experiments with model membranes referred to as large unilamellar vesicles (LUVs). The use of membrane models allowed us to modulate the lipid composition of LUVs and control the system temperature. By adding fluorescence probes, we were able to assess the mechanical properties of the membranes, particularly their lipid phases and membrane fluidity. Interestingly, membrane fluidity is related to the fluorescence “Generalized Polarization” parameter (GP) quantified with environmental fluorescent probes (see materials and methods). Ordered (rigid) membranes have high GP values and disordered (fluid) membranes have low GP values. As defined in the methods section (X – control), negative values of the delta-GP indicate an increase in fluidity and positive values indicate an increase in rigidity compared with those of the controls^43^. Since the most significant changes in sterol distribution following the incubation of cilia with the different inhibitors were observed with AY9944 (DHCR7 inhibitor), we decided to focus our study on the effect of four specific sterols (7-DHC, 8-DHC, desmosterol and cholesterol), on membrane fluidity.

We compared the fluidity of PC/SM/Chol LUVs (1/1/1 mole) with that of PC/SM/Sterol (sterol being Desmo, 7DHC, or 8DHC). As presented in Figure 4a, the membranes containing 8DHC exhibited a degree of fluidity very similar to that of the membranes containing cholesterol (corresponding to the x-axis 0 value). In contrast, a significant increase in fluidity was observed for membranes containing desmosterol and, to a lesser extent, in those containing 7DHC. These effects were observed at 25 and 37°C. Second, we investigated whether the cholesterol presence in membranes influenced these sterol properties. Therefore, we prepared PC/SM/Chol/sterol (1/1/0.5/0.5 mole) LUVs (sterol being: chol, Desmo, 7-DHC or 8-DHC). As shown in Figure 4b, effects similar to those observed in cholesterol-free membranes were detected, although with less pronounced variations. Finally, on the basis of the lipid analysis by mass spectrometry, we prepared model membranes mimicking “control” cilia (PC/SM/chol/Desmo/8DHC, 10/10/7.5/2/0.5 mole) and cilia treated with AY9944 (10/10/6/0.5/3.5 mole), with a reduction in cholesterol and desmosterol and an increase of 8DHC. Figure 4c shows that the membranes mimicking the effect of AY9944 exhibited increased rigidity (increase in GP and ΔGP) at both tested temperatures. This effect is probably due to the replacement of the fluidifying desmosterol by the more rigidifying 8-DHC, which could explain the resulting defects in Hedgehog signaling.

**Figure 4.**
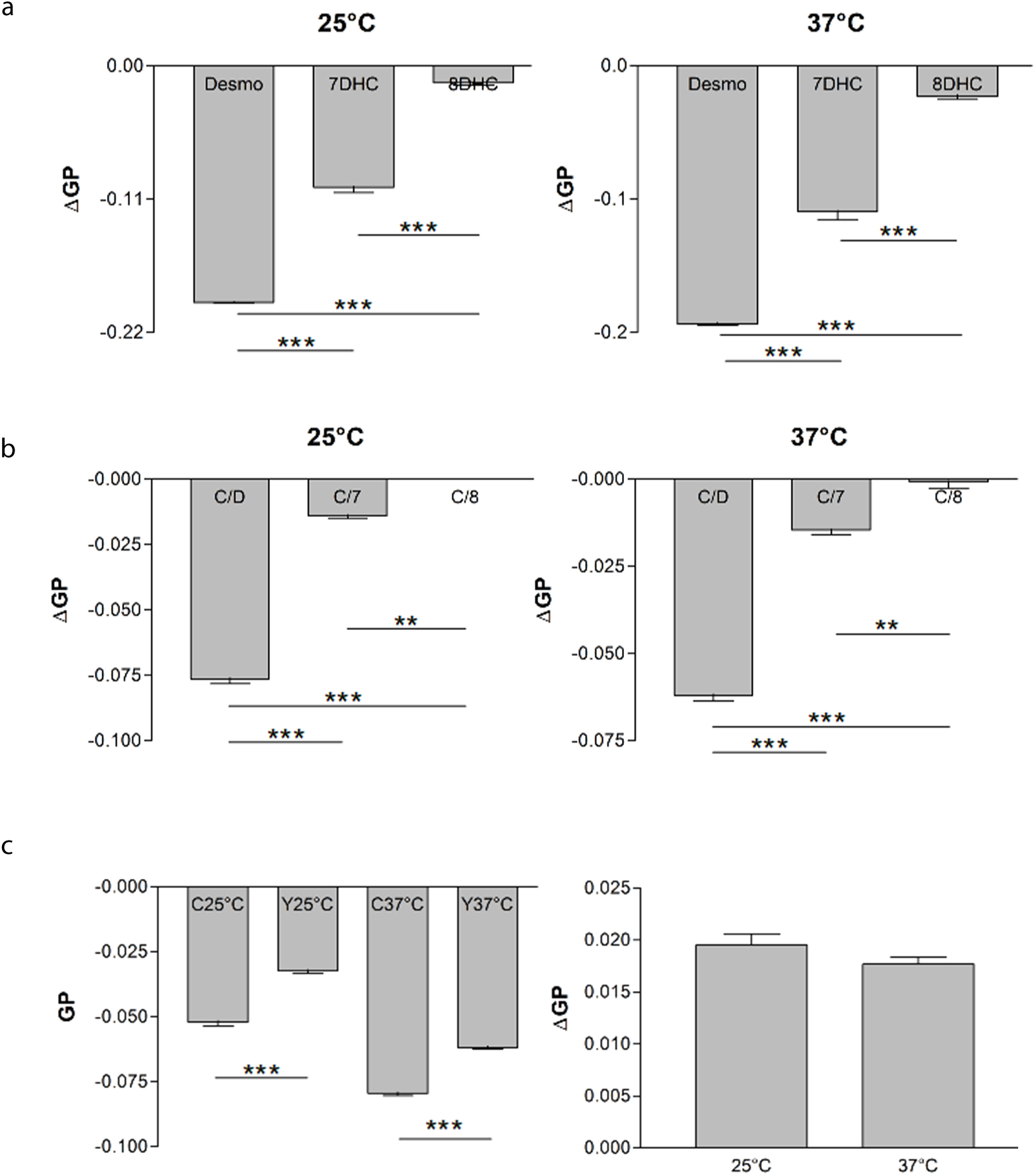
SLOS sterol composition leads to stiffer membranes. Effect of different sterols on lipid compaction (fluidity) in membrane models studied with the fluorescent probed Di-4-ANEPPDHQ. (a) Change in fluidity expressed as ΔGP change (GP_SterolX-LUV_ – GP_cholesterol-LUV_) in LUVs of different composition at 25 and 37°C. PC/SM/Desmo, PC/SM/7dhc and PC/SM/8dhc, and PC/SM/chol (as “control”) (1/1/1 mol ratio). Means ± SD of 17 and 12 experiments for 25 and 37°C respectively. (b) Change in fluidity for PC/SM/chol/Desmo, PC/SM/chol/7dhc and PC/SM/chol/8dhc (1/1/.05/0.5 mol ratio) versus PC/SM/chol. Means ± SD of 10 and 12 experiments for 25 and 37°C respectively. (c) GP values of LUVs mimicking the control “C” and the AY9944 “Y” treated cilia membranes (PC/SM/Chol/Desmo/8dhc, see text) at 25 and 37°C. (d) Change in fluidity expressed as ΔGP change (GP_AY9944-LUV_ – GP_control-LUV_) of the LUVs in (c). Means ± SD of 6 independent experiments. ** P<0.01, *** P<0.001 by student t test.

### Sterol alteration modifies ciliary length and the cell response to Sonic Hedgehog

To study the effect of sterol-content modulation in ciliary function, we measured ciliary length in treated versus untreated cells (Fig.5a). Cilia on MEFs treated with AY9944 were slightly longer on average than those of untreated cells. SAG (Smoothened Agonist) treatment resulted in slightly shorter cilia, whereas other drugs did not significantly impact ciliary length. As it is difficult to interpret minor changes in ciliary length, other than indicating that the cilium overall is affected by the treatment, we investigated Hedgehog pathway’s activation. We measured the mRNA content of Gli1, a transcription factor of the Hedgehog pathway that is upregulated upon pathway activation, via RT-qPCR. Figure 5b shows that compared with WT untreated MEFs, cells treated with any of the cholesterol modulator presented a lower concentration of Gli1 mRNA which was explained by a reduced capacity to activate the Hedgehog pathway. This effect appeared stronger with simvastatin, indicating that ciliary sterol homeostasis has a major impact on Hedgehog signaling function. This simvastatin result aligns with Maerz and colleagues findings regarding atorvastatin^12^.

**Figure 5.**
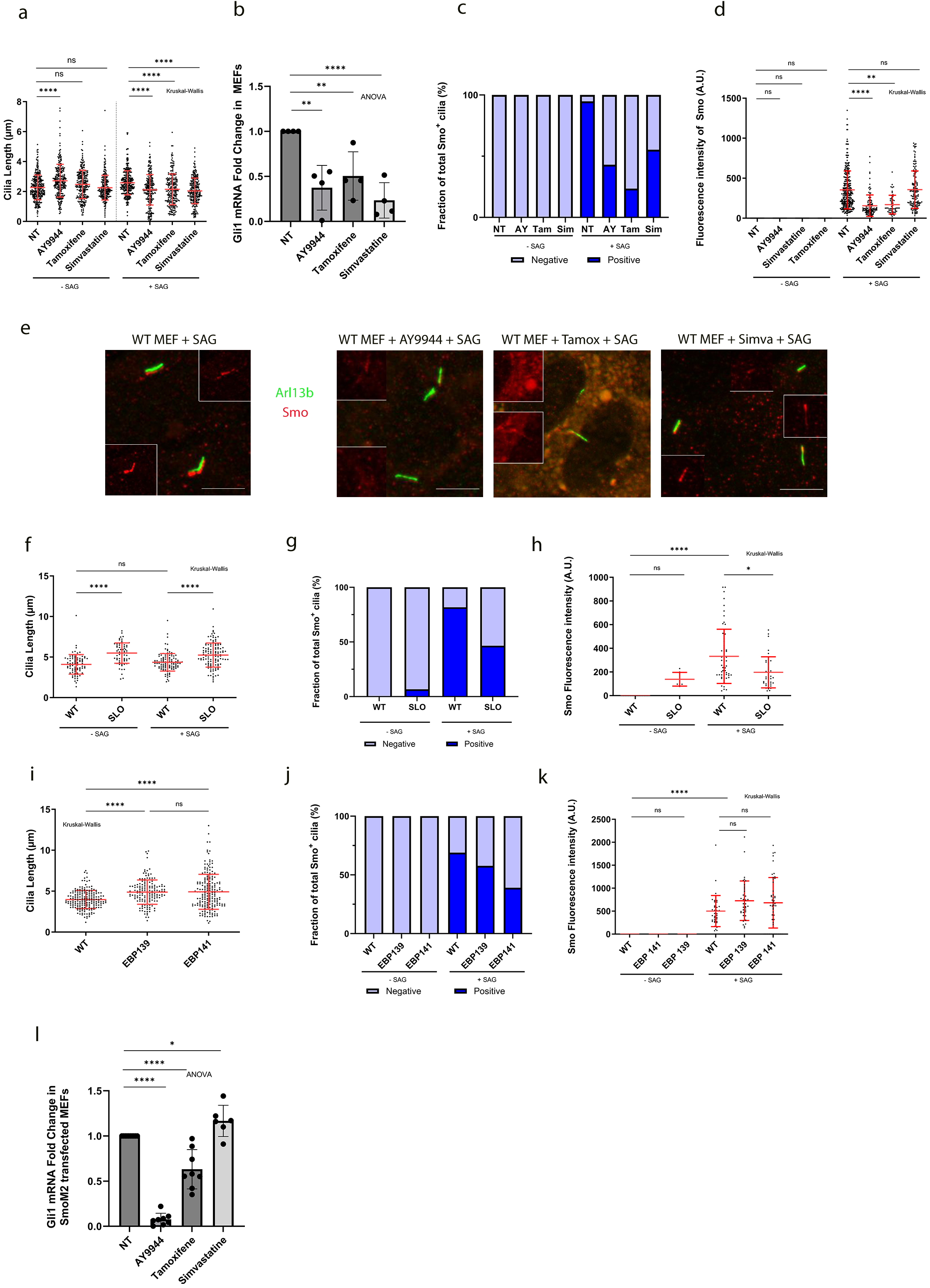
Sterolome modulation leads to Hedgehog defects. (a) Primary ciliary length in WT MEFs with (AY9944, Tamoxifen and Simvastatin) or without (NT) treatment or SAG. (b) qPCR of MEF RNA using Gli1 as a readout of Hedgehog pathway activation by SAG with (AY9944, Tamoxifen and Simvastatin) or without (NT) treatment. (c) Fraction of total Smo-positive cilia in percentage in WT MEFs with AY9944 (AY), Tamoxifen (Tam), Simvastatin (Sim) or without (NT) treatment or SAG. (d) Fluorescence intensity of Smoothened in the cilium of WT MEFs in arbitrary units (A.U) with AY9944 (AY), Tamoxifen (Tam), Simvastatin (Sim) or without (NT) treatment or SAG. (e) Immunofluorescence on WT MEFs with antibodies directed against the ciliary marker Arl13b (Green) and Smoothened (Smo, Red) after activation of the pathway by SAG with or without AY9944 (AY), Tamoxifen (Tam) and Simvastatin (Sim). Close ups show the Smoothened channel, bars = 5µm (f) Primary ciliary length in healthy (WT) and SLOS donors (SLO) fibroblasts activated by SAG. (g) Fraction of total Smo-positive cilia in percentage in healthy (WT) and SLOS donors (SLO) fibroblasts. 3 independent experiments. (h) Fluorescence intensity of Smoothened in the cilium in arbitrary unit in healthy (WT) and SLOS donors (SLO) fibroblasts activated or not by SAG. (i) Primary ciliary length in healthy (WT) and EBP CRISPR (EBP 139 and EBP141) fibroblasts. (j) Fraction of total Smo-positive cilia in percentage in healthy (WT) and EBP CRISPR (EBP 139 and EBP141) fibroblasts. 3 independent experiments. (k) Fluorescence intensity of Smoothened in the cilium in arbitrary unit in healthy (WT) and EBP CRISPR (EBP 139 and EBP141) fibroblasts activated or not by SAG. (l) qPCR of WT MEFs Gli1 RNA from cells with (AY9944, Tamoxifen and Simvastatin) or without (NT) treatment. 2 independent cell lines,. For b) and l) Means ± SD of 4 independent experiments. *p<0,01; ****p<0,0001 by ANOVA. For the rest, Means ± SD of 3 independent experiments. n.s not significant; **p<0,01, ****p<0,0001 by Kruskal Wallis.

### Ciliary sterol alteration profoundly modifies Hedgehog signaling at the level of Smo

To better understand the effects of the various pharmacological treatments on primary cilia function, we studied the activation of the pathway in more details. This step is strongly dependent on Smoothened (Smo), thus, we examined Smo localization after SAG treatment by immunofluorescence (Fig.5c-e). Under normal conditions, Smo entered the cilium upon activation as expected (Fig.5c,e). However, both AY9944 and tamoxifen-treated cells exhibited a drastic reduction in the number of Smo-positive cilia (Fig.5c,e), as well as a weaker Smo signal (Fig.5d,e) indicating that when Smo can enter the cilium, it is less retained than in untreated cells. In Simvastatin-treated cells, the number of Smo-positive cilia was also reduced, however, Smo appeared to be properly retained after entering the cilium. These results were confirmed in skin fibroblasts harvested from one SLOS patient and one healthy donor (Fig.5f-h). The findings were consistent with those obtained with AY9944 treated MEFS in terms of Smo-positive cilia (Fig.5g) and Smo fluorescence intensity (Fig.5h). It indicates that AY9944 treatment efficiently mimics the patient’s genetic defect. As we could not find a CDPX2 patient cell line with mutations in EBP, we generated two independent EBP CRISPR cell lines from healthy donor cells. After verifying that EBP expression was indeed abolished in these cells (Fig.S1a), we subjected them to the same experiment as SLOS patient cells and were able to show that results with human cells aligned with our tamoxifen experiments (Fig.5i-k), although the effect of tamoxifen on the number of Smo-positive cilia (Fig.5j) and on the cell’s ability to retain Smo appeared to be more drastic (Fig.5k), indicating that an off-target effect of the drug could explain this difference. Taken together, these results are in accordance with those of the study by Blassberg and colleagues^13^ on SLOS. The results of experiments with Tamoxifen and EBP CRISPR cells were surprising, as, unlike SLOS, CDPX2 phenotypes have not been formally linked to Hedgehog defects in the literature.

### Defects in the Hedgehog pathway are not limited to the activation steps

As cholesterol and its derivatives are extremely important in pathway activation^21,22,45,46^, we reasoned that bypassing the activation steps would be appropriate for studying the possible defects downstream of these steps and verifying the effect of altering the membrane composition, without interference from other roles cholesterol might play in the pathway. To do so, we used a constitutively active form of smoothened (SmoM2) that activates the pathway at the Smo level. Upon transfection with SmoM2, two independent stable cell lines exhibited a strong Hedgehog response as shown by Gli1 expression (Fig.S1b). When treated with AY9944 or tamoxifen, the cell line demonstrated a reduced capacity to activate the pathway showing that altering ciliary membrane sterols led to defects that were not limited to the activation steps (Fig.5l). However, the reduction was not of the same order of magnitude, as AY9944 almost completely abolished Gli1 production, whereas tamoxifen only reduced it by half. Interestingly, treatment with simvastatin did not hinder the ability of SmoM2 to activate the pathway and even had a slight propensity to promote this process. This important set of results helps explain the differences in the strength of Hedgehog defects in patients affected by different mutations and pathologies (Fig 6) and the apparent discrepancy between these defects and the overall reduction in pathway activation (Fig.5a).

**Figure 6.**
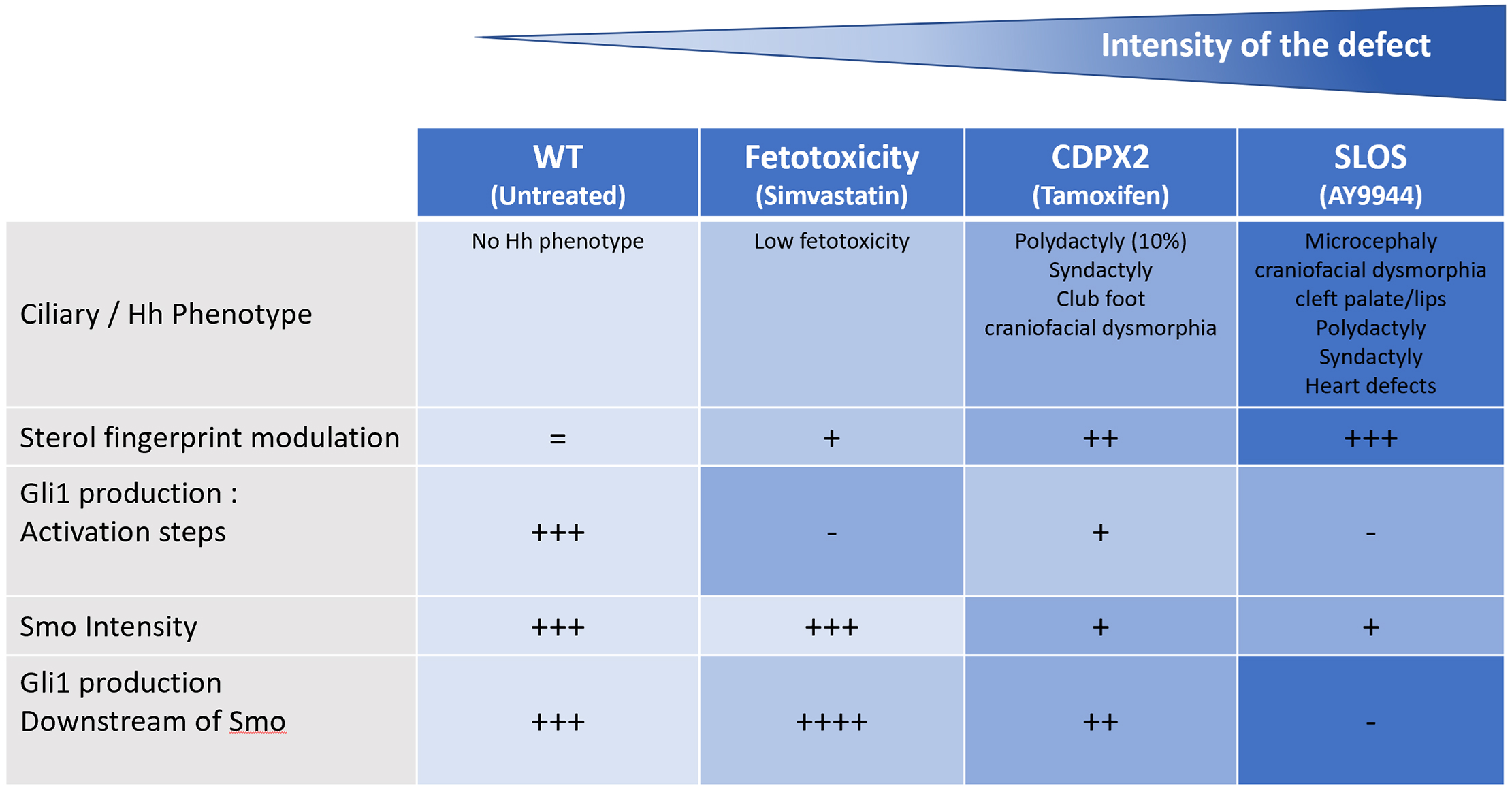
Intensity of Hh phenotypes are mirroring intensity in Hh molecular defects. Graphical summary of the intensity of Hh defects

## Discussion

Here, we provide a comprehensive overview of the sterol composition of primary ciliary membranes in mammalian cells via mass spectrometry-based metabolomics. Previous publications have characterized the lipid composition of ciliary or flagellar membranes but using less sensitive and specific analytic methods^47–50^ (see also for a review^51^). Moreover, these studies were generally performed in unicellular organisms (*e.g Paramecium and, Tetrahymena*), as they are widely known to shed their cilia by autotomy, a process that is not well conserved in mammalian cells. Additionally, most of those studies did not consider sterols specifically. Although many studies have investigated flagellar cholesterol with fillipin and electron microscopy, this method cannot differentiate various sterol species^52–54^. More recently, Lobasso and colleagues ^55^ analyzed cilia isolated from porcine olfactory epithelial cells by mass spectrometry, however, this work was not specifically focused on sterols and, olfactory cilia are mainly sensory and not primary cilia. To bridge this gap, we set up a method to isolate primary cilia from MDCK cells adapted from Ishikawa and Marshall’s work^56^. We used dibucaine for deciliation, which allowed us to recover longer and more abundant cilia from less starting material versus that achieved with the original method. The sterol fingerprint of a generic mammalian primary cilium is an important piece of data, as it enables future comparisons with cilia from various altered conditions, as shown in this work, where we modulated the cholesterol synthesis pathway. Our results clearly revealed that the sterol membrane composition is tightly regulated by the cell, as shown by the high cholesterol concentration in cilia compared to other cell membranes, as well as the inability of 7-Lathosterol to enter the primary cilium even when treatment increases its concentration in the rest of the cell, suggesting a specific sterol barrier. These sterols changes could result in the modulation of the physicochemical properties of the ciliary membrane as suggested by our fluidity experiments. More work is still needed to understand precisely how this lipid homeostasis is regulated, and whether the transition zone plays a role in this regulation.

The role of cholesterol in Hedgehog signaling has long been known because the secreted morphogen Sonic Hedgehog is modified with a cholesterol moiety adduct to ensure its activity on Patched ^21,22^. In recent years, many efforts have been made to better understand how cholesterol and its hydroxylated derivatives (oxysterols) can activate the pathway at the Smo level ^23–25,46,57–60^. Oxysterols bind to the Smo N-terminal extracellular cysteine-rich domain (CRD), leading to Smo accumulation in cilia, and subsequent pathway activation ^24,25,61–64^. Cholesterol also binds to the Smo CRD and induces Hedgehog signaling, thus, cholesterol is a plausible endogenous Hedgehog activator^23,65–67^. Two important studies recently revealed that membrane-bound cholesterol, which is organized in a small pool referred to as “accessible cholesterol”, could be the main form of cholesterol activating Smoothened ^26,27^. One caveat regarding the existing literature is that there is no acknowledgment that the membrane contains other sterols than cholesterol; however, several of other sterols produced in the cholesterol synthesis pathway demonstrate, as we observed in this study, different biophysical properties with regard to the membrane. The lack of consideration for other sterol species is illustrated by the wide use of M-β cyclodextrin ^12,13,26,27^ in experiments aimed at removing cholesterol from membranes although this agent is not a specific chelator of cholesterol and also removes other sterols. The fact that at least one disorder in the cholesterol synthesis pathway, the Smith-Lemli-Opitz Syndrome, leads to Hedgehog defects, an elevation of both 7-DHC and 8-DHC as well as a reduced cholesterol concentration, prompted us to evaluate the consequences of increased sterol intermediate concentrations in the ciliary membrane. To do so, we mimicked two molecular defects responsible for human pathologies, and used AY9944 to block the DHCR7 enzyme responsible for SLOS and tamoxifen to block the Δ7-Δ8 isomerase responsible for X-linked Dominant Chondrodysplasia Punctata 2. These two pathologies were chosen because SLOS is known to be linked to the Hedgehog pathway^12–14^, whereas no association has been reported for CDPX2. We also used simvastatin, a well-known blocker of HMG-CoA reductase, as this enzyme appears very early, before squalene formation, whereas the two other enzymes play a role in the late steps of cholesterol synthesis. The three pharmacological agents exerted very different effects on the sterol composition of the ciliary membrane, prompting us to investigate the consequences of such changes on the biophysical properties of the membrane. Using large unilamellar vesicles, we showed that modeling the sterol composition of DHCR7 blockade led to stiffer membranes. This could at least partially explain the inability of the cells to fully activate the pathway upon SAG addition due to poor Smo ciliary entry in the treated cells, which aligns with previous findings^13^. However, these experiments could not decipher between the activation steps known to be at least partially controlled by cholesterol and the rest of the pathway. Hence, we established stable cell lines expressing a constitutively active form of Smo, SmoM2, which translocates to the cilium and activates the pathway at the Smo level. These cell lines presented a major defect in pathway activation upon AY9944 treatment, indicating that membrane sterols not only play a role in Smo activation by cholesterol but are also essential for signal propagation downstream of Smo. Tamoxifen also decreased Gli1 production in SmoM2 cells, although not as drastically as AY9944. Finally, conversely to other treatments, simvastatin-treated cells produced slightly more Gli1. Taken together, these results are in agreement with the hypothesis that the biophysical state of the membrane, partially established by precise sterol concentrations, is as important to the proper function of the Hedgehog pathway as the accessible cholesterol theory is.

The results concerning CDPX2 were surprising, as we also observed a limited ability of the tamoxifen-treated cells to activate the pathway upon SAG addition, despite a lack of previous association between this pathology and Hedgehog signaling. Interestingly, in the literature concerning inborn cholesterol metabolism errors, many patients, including those with CDPX2, exhibit symptoms reminiscent of Hedgehog defects, such as polydactyly^10^. Notably, there is a gradient in the severity of Hedgehog-specific symptoms ranging from light (simvastatin)^68^, to mild (CDPX2) to severe (SLOS). This gradient parallels the severity of sterol modulation in the ciliary membrane and the severity of Hedgehog defects during activation and downstream signaling (Fig. 6). Thus, this study represents a first step toward a better understanding of the Hedgehog mechanisms underpinning inborn errors of cholesterol metabolism.

## Resource availability

Lead contact

The lead contact is Dr Thibaut Eguether

Materials availability

Plasmids generated during this study are available upon request.

Data and code availability

Unprocessed data files generated during this study are available upon request.

## Acknowledgements

We would like to thank Drs Paul Guichard and Anne-Marie Tassin for their careful reading of the manuscript.

## Authors contributions

Conceptualization, T.E and A.L.; Methodology, T.E,C.D and S.M, Investigation T.E, L.K, S.M, J.A.S, N.B, M.F, G.D and C.D, Formal Analysis T.E, L.K, G.D and A.C, Writing – Original Draft, T.E and J.A.S., Writing –Review & Editing, T.E, A.L, L.K., G.J.P and J.A.S; Funding Acquisition, T.E, G.T and A.L; Resources, T.E, G.J.P, T.B and B.I; Supervision, T.E.

## Supplemental information

Figures S1

---

**Figure S1:**
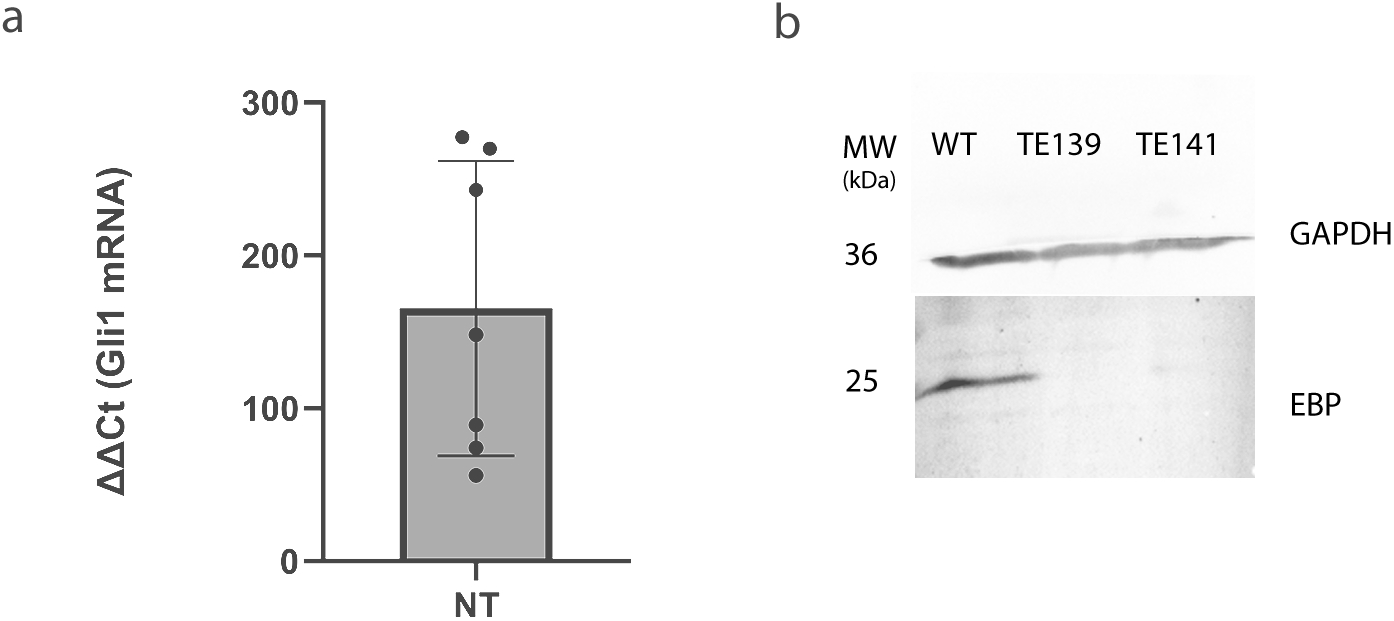
(a) Western Blot of Healthy donors fibroblasts (WT) and EBP CRISPR cells (EBP 139 and EBP141). Top : GAPDH antibody. Bottom : EBP antibody.(b) qPCR of MEF RNA. Gli1 response to SmoM2 transfection in WT MEFs. Two independent cell lines, 3 independent experiments.

